# dbSUPER: a database of super-enhancers in mouse and human genome

**DOI:** 10.1101/014803

**Authors:** Aziz Khan, Xuegong Zhang

## Abstract

Super-enhancers are the clusters of transcriptional enhancers that can drive cell-type-specific gene expression and also crucial in cell identity. Many disease-associated sequence variations are enriched in super-enhancer regions of disease-relevant cell types. Thus, super-enhancers can be used as potential biomarkers for disease diagnosis and therapeutics. Current studies have identified super-enhancers in more than 100 cell types and demonstrated their functional importance. However, no centralized resource to integrate all these findings is available yet. We developed dbSUPER (http://bioinfo.au.tsinghua.edu.cn/dbsuper/), the first integrated and interactive database of super-enhancers, with the primary goal of providing a resource for assistance in further studies related to transcriptional control of cell identity and disease. dbSUPER provides a responsive and user-friendly web interface to facilitate efficient and comprehensive search and browsing. The data can be easily sent to Galaxy instances, GREAT and Cistrome web servers for downstream analysis, and can be visualized in UCSC genome browser while custom tracks added automatically. The data can be downloaded and exported in variety of formats. Further, dbSUPER lists genes associated with the super-enhancers and links to various other databases such as GeneCards, UniProt and Entrez. dbSUPER also provides an overlap analysis tool, to annotate user defined regions. We believe dbSUPER is a valuable resource for the biologists and genetic research communities.

## INTRODUCTION

Enhancers are cis-regulatory elements in DNA that enhance the transcription of target genes by communicating with core promoter through several mechanisms including looping, tracking, linking, and relocation models (1–6). These enhancer elements contain binding sites for sequence-specific transcription factors (TFs), which help in recruiting coactivators and RNA polymerase II at target genes (2–5). Since the discovery of the first enhancer in animal virus SV40 (7), there has been tremendous development of technology and methodology to study the role of enhancers in gene expression. The number of active enhancers operating in a single mammalian cell can be in thousands while in the human genome their number can be up to ∼1 million (8, 9). Many approaches were applied for genome-wide identification of enhancers, for example chromatin immunoprecipitation followed by high throughput sequencing (ChIP-seq) for coactivator protein p300 (10), and histone modifications including H3K4me1 and H3K27ac (11, 12). H3K27ac was used as enhancer mark to identify enhancers in hESC (13) and mESC (14), and further these enhancers were grouped into active and poised enhancers.

Recently a small set of enhancers, spanning large regions of the genome in a clustered manner which are occupied by high levels of Mediator (Med1), master transcription factors and coactivators are named as super-enhancers (SEs) (15, 16). These super-enhancers drive cell-type-specific gene expression programs and many disease-associated sequence variation are preferentially enriched in these regions of disease-relevant cell types (16, 17). In cancer cells, super-enhancers are associated with different oncogenes, including MYC (16, 17). These super-enhancers were discovered through ChIP-seq experiments for the master transcription factors and Mediator (Med1), BRD4 and H3K27ac (15–17). A parallel study observed similar patterns through integrated analysis of human pancreatic islets data with nine cell types from ENCODE, and were named as “stretch enhancers” which are larger than 3kb regions (18). Downstream computational and *in vivo* analysis revealed that these stretch enhancer regions are key chromatin features for cell type-specific gene expression programs, and a sequence variation in these stretch enhancer regions promotes the risk of common human diseases (18).

Since the discovery of super-enhancers, there has been various efforts to demonstrate the functional importance of these regulatory regions in gene expression and disease. Recent studies used genome editing technique CRISPR (clustered regularly interspaced short palindromic repeats) to validate the importance of super-enhancers and their constituents, especially those associated with cell identity genes (19–21). In chronic myeloid leukemia a super-enhancer associated with GATA2 gene is responsible for 80% of GATA2 expression (19). In ESC a 13kb long super-enhancer associated with Sox2 gene is responsible for more than 90% of Sox2 expression (20). In ESC most (12/14) super-enhancer constituents led to reduced expression of the associated gene (21). Another study demonstrated that hotspots of transcription factors in the early phase of adipogenesis are highly enriched in super-enhancer regions, which drive adipogenic-specific gene expression (22). Wang *et al*., found a large number of dynamic NOTCH1 (a master regulatory protein) sites in the super-enhancer regions (23). They observed that 83% of NOTCH1 sites overlap with H3K27ac ChIP-seq peaks and demonstrated the importance of Notch super-enhancer interaction in gene expression (23). Another study identified super-enhancers by profiling BRD4 ChIP-seq signal in multiple cancer cells and found considerable loss of BRD4 at super-enhancer regions by treating cancer cells with the BET-bromodomain inhibiter JQ1 (16). Recently, a large scale collaborative research revealed highly asymmetric binding of BRD4 at super-enhancers in Diffuse large B-cell lymphoma (DLBCL) cells, and revealed that the genes regulated by super-enhancers are particularly sensitive to JQ1 inhibition (24). A significant decrease in the growth of DLBCL cells was observed after JQ1 treatment which was engrafted in mice and improved survival of mice (25). Plutzky et al extended the current understanding of super-enhancer function by discovering that super-enhancers can perform as fast switches to enable the rapid cell state transition (26, 27). Kwiatkowski et al discovered that the transcription-targeting drug THZ1, a CDK7 inhibitor, preferentially reduced the expression of genes associated with super-enhancers (28). A follow-up *in vivo* study in small cell lung cancers (SCLC) showed the association of super-enhancers with proto-oncogenes and SCLC identity genes, and transcription-targeting drug THZ1 preferentially targets super-enhancer-driven genes (29, 30). Mansour et al extended the functional importance of super-enhancers by showing that somatic mutations introduce binding sites for MYB transcription factor, which creates a powerful super-enhancer that mediates the overexpression of oncogenes in T-cell acute lymphoblastic leukaemia (T-ALL) (31, 32). Later, studies have linked the activation-induced deaminase (AID) off-targeting activity to the process of convergent transcription (33) and these AID targets are mainly grouped within super-enhancer and regulatory clusters (34). Very recently, Vahedi et al found super-enhancers in T-cells by profiling p300 ChIP-seq signal and showed disproportionate alteration in the expression of rheumatoid arthritis risk genes with super-enhancer structures by treating T-cells with the Janus kinase (JAK) inhibitor tofacitinib (35). Also, disease-associated SNPs for autoimmune diseases, including rheumatoid arthritis, were highly enriched in super-enhancer regions (35). The current research effectively demonstrates the importance and potential application of super-enhancers as they can play key roles in cell identity and diseases. The concept of super-enhancer is still evolving but has already gained extensive attention in the community. More systematic and comprehensive studies will lead to more accurate understanding of the concept as well as its functions.

The above mentioned studies have generated large amount of data by profiling ChIP-seq signal for Mediator complex (Med1), master transcription factors (MyoD, T-bet and C/EBPα) (15), H3K27ac (17, 24, 29), BRD4 (16, 24) and p300 (35) in different tissue and cell types. The data produced by most of these studies is shared with public in the literature. But a centralized database to integrate existing and forthcoming data is needed, to streamline the downstream analysis and to answer many questions related to these newly discovered regions. Hence, we developed a user-friendly and interactive database of super-enhancers by integrating all the published and new data with the aim to provide a resource to help bioinformaticians and biologists to perform further analysis and study of transcriptional control of cell identity. We named the database dbSUPER which is available for academic use at http://bioinfo.au.tsinghua.edu.cn/dbsuper/. The database can help the research community to search, browse, export, send and download super-enhancer-related data in more systematic way. The database will be updated with latest progresses in the field and we hope it will be a helpful tool to better enable downstream analysis of super-enhancers and their role in gene regulation.

## MATERIAL AND METHODS

### Data sources

The current version of dbSUPER contains data collected from a variety of sources and also produced by using the published pipeline (17). We stored this data into a MySQL based database after preprocessing for fast and efficient query. We collected 2,558 super-enhancer regions for 5 mouse tissue and cell types including mESC, pro-B cells, myotubes, Th cells, and macrophages in the mouse genome (15). These super-enhancers were identified by ranking ChIP-seq signals for Med1 for mESC and pro-B cells, and MyoD, T-bet and C/EBPα for myotubes, Th cells, and macrophages respectively (15). We collected 58,283 super-enhancer regions for 86 human tissue and cell types in the human genome, which were identified using H3K27ac ChIP-seq signal based ranking (17). We collected super-enhancers for three cells in small cell lung cancer (SCLC) including NCI-H69, GLC16 and NCI-H82, which were identified by profiling ChIP-seq signal for H3K27ac (29). We collected super-enhancers for six cells in Diffuse Large B Cell Lymphoma (Ly1, DHL6, Ly3, HBL1, Ly4, and Toledo) and one human tonsil, which were identified by profiling ChIP-seq signal for H3K27ac (24). We also identified 2,475 super-enhancers in NHEK, HSMM and pancreatic islets cells using published pipeline (17). In total, current version of database contains 68,508 super-enhancers (mean size of 33,660bp and average number of 659 super-enhancers in each cell-type) in 99 human and 5 mouse tissue/cell types. A detailed list of all tissue/cell-types including number of super-enhances, mean size (bp) and identification method used for each can be found in Supplementary Table S1.

### Computational methods for super-enhancer identification

Besides the data from the above references, we identified super-enhancers in human cells NHEK, HSMM and pancreatic islets using ChIP-seq data for H3K27ac, as described in (17). An overview of super-enhancer identification and data integration is presented in Figure 1. Initially, ChIP-seq data for H3K27ac was downloaded from GEO and mapped to reference genome (hg19) using Bowtie (version 0.12.9) (36). MACS (version 1.4.1) was used to identify enriched regions as enhancers with a threshold (p value 1 x 10^-9^) (37). ROSE (Rank Ordering of Super-Enhancers) algorithm (https://bitbucket.org/young_computation/rose) was used to separate super-enhancers from enhancers (15, 16). We used 12.5kb distance threshold to stitch enhancers together. Then these stitched enhancers were ranked, based on the ChIP-seq occupancy of H3K27ac, which revealed a geometrical inflection point and established a cut off that separates super-enhancers from typical enhancers. An implementation of ROSE algorithm and more detailed definition of super-enhancer concept can be found elsewhere (15, 16, 38). The current version of database is developed using hg19 assembly for human genome and mm9 assembly for mouse genome. For any of the data which was not available in this assembly, we adjusted the coordinates using UCSC genome browser using liftOver tool (39).

**Figure 1.**
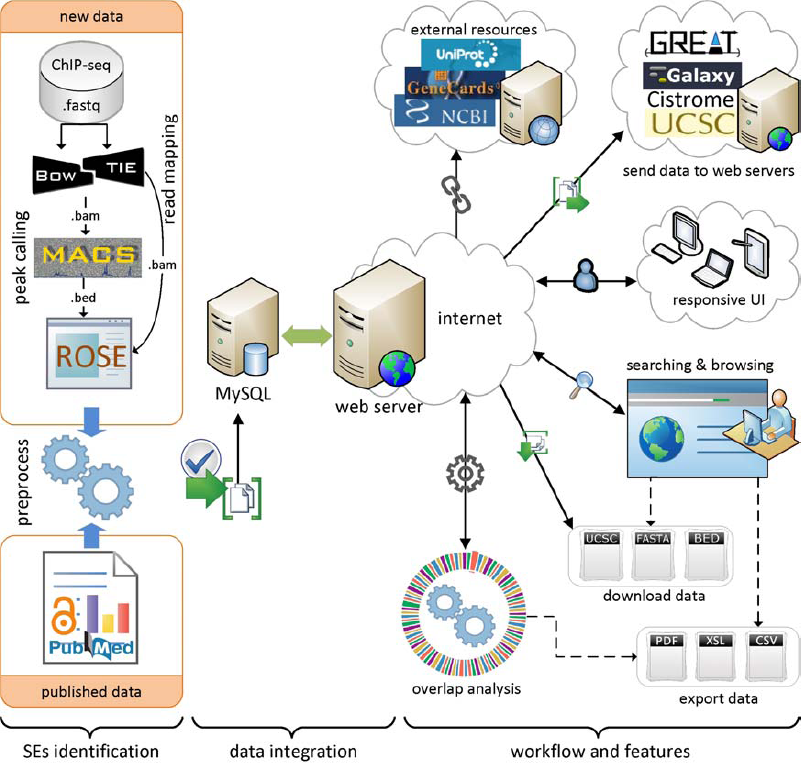
Overview of super-enhancer identification, data integration, and dbSUPER workflow and features.

ChIP-seq data for various chromatin regulators and coactivators including p300 (10) is used for enhancer identification. But in the case of super-enhancers, which are separated from a list enhancers based on ChIP-seq occupancy for certain factors including Mediator (Med1). Initially, super-enhancers were identified using Med1 ChIP-seq signal (15), but using H3K27ac (17) and BRD4 (16) we can achieve comparable results. A list of all the ChIP-seq data with GEO accession for data collected and generated can be found on “Data Sources” page. So far different ChIP-seq-based ranking methods have been used to identify super-enhancers but still a conceptually appealing definition and a set of functionally important features yet to come (38).

### Assigning genes to super-enhancers

dbSUPER also contain associated genes for all the super-enhancers. These transcriptionally active genes were assigned to super-enhancers using a simple proximity rule. It is known that enhancers tend to loop and associate with target genes in order to activate their transcription (40), while most of these interactions occur within a distance of ∼50kb of the enhancer (41). Hence, using a distance threshold of 50kb, all transcriptionally active genes (TSSs) are assigned to super-enhancers within a 50kb window. This approach identified a large proportion of true enhancer/promoter interactions in embryonic stem cells (42).

## DATABASE FEATURES

### General web interface and database

The web interface of dbSUPER provides an interactive solution for searching, browsing, visualizing, downloading, exporting and transferring the data to other public servers. To access all these features, the interface provides a navigation bar on left side and footer. A quick search box is available on the home page, which can be used for fast searching and browsing. The database provides an advanced search feature to filter the super-enhancers based on more detailed criteria. For organized browsing, dbSUPER displays data in paginated, sortable and responsive tables. The responsive feature allows changing table shape to fit the data into a screen based on the user’s device resolution by adding a plus sign at the beginning of each row. By clicking the super-enhancer ID, users can view general details, details about associated genes, FASTA sequences and also links to external sources including UCSC (43), NCBI RefSeq (44) and Entrez Gene (45), GeneCards (46), UniProt (47) and Wikipedia. The data can be downloaded in different formats including BED, FASTA and UCSC custom tracks. The user-queried data can be exported to Excel, CSV and PDF files and copied to clipboard. To make the downstream analysis faster and efficient, the dbSUPER interface provides links with any Galaxy (48) instance, GREAT (49) and Cistrome (50) server to send data with one click. The interface also provides a link to visualize data in UCSC genome browser (39) by adding custom tracks automatically. The overlap analysis tool allows users to check the overlap of the regions of interest with the current database and outputs the overlapped regions in a responsive table. Further, dbSUPER plots the distribution of overlapped regions with each cell/tissue type while the overlapped regions can be exported to different formats. In the following sections, we will explain these features in detail. Figure 1 illustrates the general workflow, features and user-interface of the database.

### Searching and browsing

dbSUPER supports comprehensive user-friendly searches in different ways to bring the data to users in a more productive way. Figure 2 illustrates the interactive searching and browsing activity of dbSUPER. The home page of the website provides a quick search utility which can be used to query the database for genes of interest, cell/tissue types and enhancer identification marks. This search uses jQuery-based auto-completion features to help and guide user to discover vocabulary available in the database. After clicking the search button, a new page will display the queried data in a responsive table. The database can be browsed for each tissue or cell-type by clicking the “Browse Database” tab on the left-side navigation menu. An “Advanced Search” link is available as an option for more detailed search. The results will be displayed in a dynamic tabular form with sorting and filtering options. In this table (Figure 2C) each row is a super-enhancer and each column contains region-specific information including; ID maintained by our database, genomic loci, size, associated gene, method used to rank enhancers, rank of super-enhancer, cell/tissue type, genome and a link to UCSC genome browser. If user browses the database on a mobile device such as smartphones or tablets, a “+” sign will appear at the beginning of each row for hiding some information. The hidden information will not be displayed unless user touches the “+” sign, as shown in Figure 2E. This will avoid horizontal scrolling, by making data fit into the screen. By default, each page displays 25 records and user can view remaining records using the pagination features on the bottom right of the table. The number of records in each page can be increased / decreased between 10, 25, 50 and 100 using the “records per page” dropdown menu. The tabular data can be filtered using the search box on the top right and the user can sort the data based on any field of interest. Details about each super-enhancer can be viewed by clicking the super-enhancer ID. Beside the general details about the super-enhancer, it also lists details about the associated gene, which includes information like gene symbol, chromosome, transcription start site, transcription end site, strand and number of exons for the gene. Further, dbSUPER provides external links to UCSC Genome Browser (43) (http://genome.ucsc.edu/), NCBI Gene (44, 45) (http://www.ncbi.nlm.nih.gov/gene/), GeneCards (46) (http://www.genecards.org), UniProt (47) (http://uniprot.org) and Wikipedia (http://en.wikipedia.org) to facilitate users to use those sources to study the selected super-enhancer on the corresponding aspects. For each super-enhancer region, a FASTA sequence file can be viewed and downloaded.

**Figure 2.**
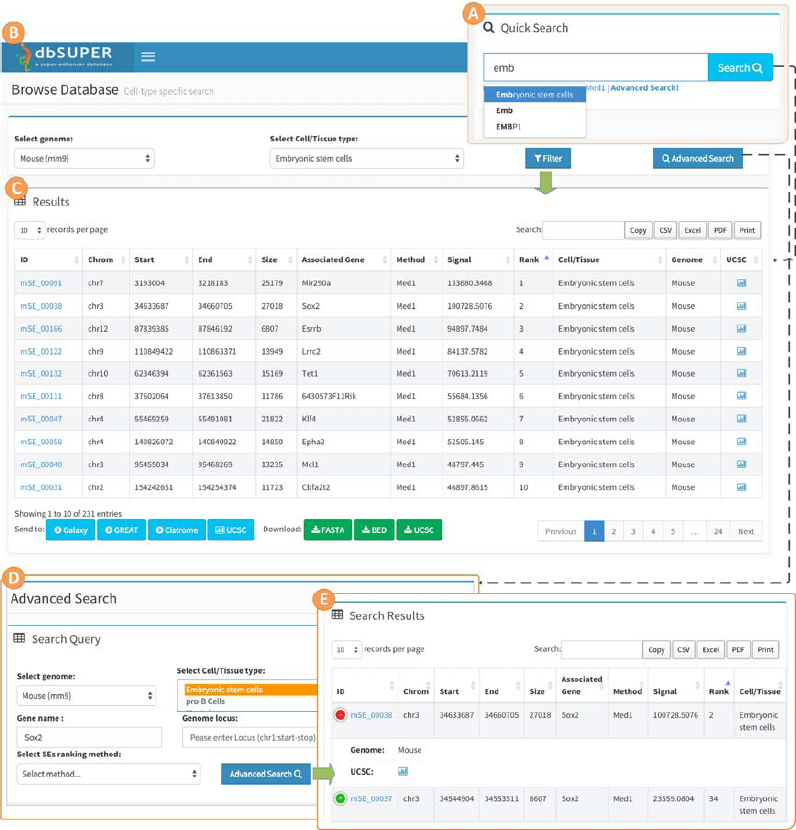
Interactive searching and browsing activity of dbSUPER. (A) Quick search feature on the homepage. (B) Browse database for mESC cell-type. (C) Results for browse database and also for quick search using keyword “mouse embryonic stem cell”. (D) Advanced search page, look for Sox2 gene in mESC. (E) Results of advanced search query.

### Data download and export

We provide all the data in multiple formats including BED, FASTA and UCSC custom tracks for users to download. Downloading can be performed either from the download page or during browsing cell/tissue specific data in the browse section. The user-queried data can be exported as Excel, CSV and PDF files, using the respective buttons on the top right of each data table. dbSUPER also provides the one-click feature to copy data tables to the clipboard and also printing features. The data can be provided freely in rational files upon request.

### Linking with other web servers and visualization

In order to provide a one-stop solution for searching and facilitating the downstream analysis including functional annotation and visualization, we provide features to directly transfer data from our database to external web servers without downloading. Super-enhancers for one or more than one cell-type or sample can be send and visualized at once. Currently, dbSUPER supports any public/personal instance of Galaxy (48) including Cistrome (50), and GREAT (49) to which data can be send directly. These features can be found under “Visualize and Send Data” tab of the user-interface. The Supplementary Figure S1 shows a demo run for cell-type LNCaP.

#### Linking with Galaxy server

Galaxy is a very useful public web server, which can be used for intensive data analysis using many integrated tools, creating pipelines, storing data and sharing analyses with others. dbSUPER provides a handy facility to directly send data for each cell/tissue type or a collection of cell/tissue type to more than sixty Galaxy (48) instances for downstream analysis. A list of publically available instances of Galaxy server can be found at https://wiki.galaxyproject.org/PublicGalaxyServers. Some of these Galaxy instances require users to register to perform the analysis, so users need to register and login before loading data from our database. To send data for a single sample user can click the Galaxy logo next to cell/tissue type of interest, dbSUPER will add a BED file to Galaxy history. To send data for more than one cell/tissue type together, user need to select the cell/tissue types of interest first and click the respective button at the bottom of table. For users who wish to send data to other public/personal Galaxy instance, the interface provides a facility to change the URL of default Galaxy server (https://usegalaxy.org). After changing the default URL, user can send the data using procedure mentioned above.

#### Linking with GREAT server

To perform functional prediction of super-enhancers by analysing the GO (Gene Ontology) annotations of the nearby genes and assigning biological meaning to them, we linked dbSUPER with GREAT (49) web server and provided a one-click facility to load data from our database to GREAT, and to automatically perform analysis. This feature can also be found under “Visualize and Send Data” tab of the user-interface and by clicking the GREAT logo next to cell/tissue type of interest.

#### Linking with Cistrome server

dbSUPER provide features to send data directly to Cistrome analysis pipeline (50) to perform correlation analyses, gene expression analyses and motif discovery.

Currently, Cistrome requires users to register to perform the analysis, so users need to register and login at (http://cistrome.org/ap/) before loading data from our database to Cistrome.

#### Visualizing in UCSC genome browser

A single super-enhancer region or super-enhancers of different cell/tissue type can be visualized in UCSC Genome Browser (43). This feature can be found on the “Visualize and Send Data” page, and also the browse page of the dbSUPER user-interface. Once a user clicks the visualization icon, dbSUPER will redirect user to UCSC Genome Browser and a custom track will be added to the UCSC genome browser session automatically.

### Overlap analysis tool

We provide an overlap analysis tool to annotate user-submitted regions with the super-enhancers available in dbSUPER. We used the intersectBed tool from the BEDTools suite (51) to find overlapped super-enhancers from dbSUPER with the submitted regions. The user is required to define a minimum percentage of overlap before running the analysis. By default, a super-enhancer in dbSUPER must overlap with user defined regions by at least 10% to be reported as an overlapping super-enhancer. User can also define the minimum percentage of overlap on both the dbSUPER and the regions uploaded. The overlap analysis can be performed by clicking the “Overlap Analysis” tab and uploading regions of interests in BED format. The BED file should be in tab-delimited format without a header. After submission, the user will receive an email with a private link to the overlap analysis results. It may take a while to get the analysis results depending on the number of regions uploaded. Two donut plots will be generated: one for the ratio of overlap within the individual tissue/cell type, and the other plot will show a total overlap map. All the overlapped regions can be downloaded in BED format and also displayed in tabular form, which further adds features to export these regions as CSV, Excel and PDF files. The Supplementary Figure S2 shows an output of overlap analysis tool with necessary steps.

## TECHNICAL BACKGROUND

The current version of dbSUPER is developed using MySQL 5.5 (http://www.mysql.com) and it runs on Linux-based Apache server. We used PHP 5.3 (http://www.php.net/) for server-side scripting. The interactive and responsive user interface was designed and built using Bootstrap 3 (http://www.getbootstrap.com), a popular responsive development framework including HTML, CSS, and JavaScript. The user-interface is responsive, this means the web interface detects the user device and changes its structure and shape according to the device resolution, in order to optimize the data view. This feature makes the interface compatible across variety of devices and browsers with different screen resolution. The database can be browsed and searched using a variety of devices including smartphones or tablets. Although we recommend Google Chrome, Firefox and Safari web browsers for best results, but the database also supports other latest standard web browsers including IE version 8 and greater. We aim to improve the accessibility and user interactivity of dbSUPER by asking for user feedbacks through the contact page on our website. We are also anonymously tracking user interactions with our website including clicks, browser and device information. This will help us to know which part of our database is most important and which part needs to be improved based on user’s interactions.

## AVAILABILITY

The dbSUPER database is freely available for the research community using the web link (http://bioinfo.au.tsinghua.edu.cn/dbsuper). The users are not required to register or login to access any feature available in the database.

## DISCUSSION

Super-enhancers are cell-type specific and are associated with the key genes that drive cell-type-specific expression, and are linked to biological processes which define the cell identity. We integrated the information of these regions and their associated genes in the dbSUPER database. dbSUPER provides a rich collection of features including (i) Fast searching, browsing and visualization (ii) Downloading and exporting data in different formats including BED, FASTA, UCSC custom tracks and CSV, Excel, PDF files; (iii) Linking with external web servers including Galaxy, GREAT, and Cistrome and sending data directly to perform downstream analyses; (iv) Providing the associated genes with links to various databases including GeneCards, UniProt and Entrez, and (v) An overlap analysis tool to check the overlap of user submitted regions with dbSUPER. The overall goal of this database is to provide a comprehensive resource and a set of interactive analysis tools to facilitate the further study of super-enhancers and their functions. The responsive user-friendly web interface facilitates efficient and comprehensive searching and browsing of the data. While there are still many unclear questions on the concept of super-enhancer and even controversy in its exact molecular definition, such an organized collection of all existing data in one compact database provides researchers a handy platform for studying those questions.

Currently, dbSUPER contains 68,508 super-enhancers for 99 human and 5 mouse tissue/cell types. The current understanding and research on super-enhancers is progressing rapidly. We will keep adding more data to the database once they are available. In the future, we are interested in collecting published research on *in vivo* validation of the computationally defined super-enhancers. We hope that, as more cell-type-specific validated data becomes available, we can construct highly reliable supervised predictive models for super-enhancers. Currently, we are working on adding more features such as motif analysis, SNPs (Single Nucleotide Polymorphisms) information, tissue-specificity analysis and the use of additional datasets to find super-enhancers for other cell/tissue types. Those features will extend the value of the database. More powerful user and session management modules are also under consideration, which will enable users to save their results and sessions and share with their collaborators or the community. Based on the current progress in the field, we believe that dbSUPER will be of particular interests to people working on the molecular and systems biology of cancer and other diseases.

## SUPPLEMENTARY DATA

Supplementary Data are available at online.

## ACKNOWLEDGEMENTS

We would like to thank Dr. Richard Young at Whitehead Institute for Biomedical Research, MIT, for his useful suggestions and comments about the database and also sharing the data produced by his research lab, and thank for the anonymous reviewers for their helpful suggestions.

